# A nuclear role for ARGONAUTE-2 in regulation of neuronal alternative polyadenylation

**DOI:** 10.1101/2020.12.15.422806

**Authors:** Revital Ravid, Aviad Siany, Natalia Rivkin, Chen Eitan, Hagai Marmor-Kollet, Eran Yanowski, Alon Savidor, Yishai Levin, Gregor Rot, Gunter Meister, Eran Hornstein

**Affiliations:** Department of Molecular Genetics, Weizmann Institute of Science, Rehovot, Israel; de Botton Institute for Protein Profiling, The Nancy and Stephen Grand Israel National Center for Personalized Medicine, Weizmann Institute of Science, Rehovot, Israel; Institute of Molecular Life Sciences of the University of Zurich and Swiss Institute of Bioinformatics, Zurich, Switzerland; Regensburg Center for Biochemistry (RCB), Laboratory for RNA Biology, University of Regensburg, Regensburg, Germany

**Author notes:** Correspondence should be addressed to: Eran Hornstein, Eran Hornstein, PhD, MD., Department of Molecular Genetics, Weizmann Institute of Science, Herzl Street 1, 76100 Rehovot, Israel.

## Abstract

Argonaute 2 (AGO2), the effector protein partner of microRNAs (miRNAs) in the cytoplasmic RNA induced silencing complex, is further involved in nuclear RNA processing. However, a role for AGO2 in regulation of alternative polyadenylation was not yet demonstrated. Here, we reveal unexpected abundance of AGO2 in mouse neuronal nuclei and characterize nuclear AGO2 interactors by mass spectrometry. We discover that AGO2 broadly regulated alternative polyadenylation (APA) in neuronal cells. Specifically, we demonstrate how two isoforms of *Ret* mRNA, which encodes a receptor tyrosine kinase are regulated by AGO2-depenent APA, affecting downstream GDNF signaling in primary motor neurons.

## Introduction

Argonaute (AGO) proteins are the direct binding partners of small RNAs and primarily execute small-RNA-guided gene-silencing processes (Bartel, 2004; Hammond et al., 2001; Hock & Meister, 2008; Liu et al., 2004; Meister, 2013; Meister & Tuschl, 2004; Tuschl et al., 1999; Zamore et al., 2000). AGO2 is the most abundant among the four mammalian AGO proteins (AGO1-4), facilitating regulatory activities, based on interacting protein co-factors. GW protein trinucleotide repeat-containing gene 6 (TNRC6 A/B/C) are the chief co-factors of AGO2 that are necessary for regulatory activities, such as mRNA deadenylation, decapping and degradation (Eulalio et al., 2008; Fabian et al., 2011; Jakymiw et al., 2005; Liu, Rivas, et al., 2005; Rehwinkel et al., 2005; Zekri et al., 2009). Under normal conditions, AGO2 is primarily in the cytoplasm, in both diffuse form and in cytoplasmic foci termed processing bodies (P-bodies) (Eulalio et al., 2007; Liu, Valencia-Sanchez, et al., 2005; Schraivogel et al., 2015; Sen & Blau, 2005).

However, the presence of AGO proteins was reported also in the nucleus (Bottini et al., 2017; Chu et al., 2010; Gagnon et al., 2014; Janowski et al., 2006; Kalantari et al., 2016; Robb et al., 2005; Rudel et al., 2008; Sarshad et al., 2018; Wei et al., 2014), where they are involved in nuclear microRNA (miRNA)-mediated gene-silencing (Meister, 2013; Sarshad et al., 2018), transcriptional silencing (Benhamed et al., 2012; Chu et al., 2010; Janowski et al., 2006; Janowski et al., 2007) and splicing (Ameyar-Zazoua et al., 2012). The diversified aspects of RNA regulation (reviewed in (Nussbacher et al., 2019)), highlights the flexibility of AGO2 that is based on the specific activities of the associated RNA-binding protein cofactors that are recruited by AGO and TNRC6.

Alternative polyadenylation (APA) is a nuclear processing step of the mRNA *3* end that is executed by a nuclear complex of CPSF and CstF proteins (Danckwardt et al., 2008; Elkon et al., 2013). A canonical polyadenylation sequence is positioned in *cis* 15-30bp upstream of the polyadenylation site (polyadenylation signal, PAS), and is composed of an AAUAAA consensus sequence or variants thereof (MacDonald & Redondo, 2002; Tian & Graber, 2012). APA affects various aspects of RNA metabolism, mRNA stability, nuclear export, cellular localization and translation efficiency. Furthermore, 3’UTR isoforms directly affect the presence of miRNA recognition sites (Danckwardt et al., 2008; Elkon et al., 2013; Fu et al., 2018; Tian & Manley, 2017).

The secreted glial cell derived neurotrophic factor (GDNF) drives neurotrophic signaling that is important for axonal outgrowth, synapse maturation and neuron survival (Airaksinen & Saarma, 2002; Baudet et al., 2008; Bonanomi et al., 2012; Enomoto et al., 2001; Honma et al., 2010; Kramer et al., 2006; Pachnis et al., 1993; Runeberg-Roos & Saarma, 2007; Tuttle et al., 2019). Ret proto-oncogene is a tyrosine kinase receptor for GDNF (Romei et al., 2016) that is highly expressed in motor neurons (Baudet et al., 2008; Cintron-Colon et al., 2020; Pachnis et al., 1993). GDNF stimulation triggers RET autophosphorylation on distinct tyrosine residues, and intracellular activation of phosphatidylinositol 3-kinase (PI3-kinase) and mitogen-activated protein kinase (MAPK) pathways, which contribute to neuronal survival (Airaksinen & Saarma, 2002; Kaplan & Miller, 2000). Two RET isoforms RET9 and RET51, differ in the C-terminus sequence (Ibanez, 2013; Rossel et al., 1997). Both shorter RET9 and longer RET51 are phosphorylated on Tyr1062, but an additional tyrosine residue at position 1096, is present only at RET51 and can drive activation of GRB2 (Airaksinen & Saarma, 2002; Ibanez, 2013). The two isoforms differ in expression, intracellular trafficking and stability Consequently, RET9 and RET51, convey distinct signaling properties (de Graaff et al., 2001; Heanue & Pachnis, 2008; Lee et al., 2003; Richardson et al., 2012; Tsui & Pierchala, 2010; Wong et al., 2005). Interestingly, RET is differentially expressed and phosphorylated in amyotrophic lateral sclerosis (ALS) models (Kramer & Liss, 2015; Ryu et al., 2011; Zhang & Huang, 2006), highlighting a biomedical interest in elucidating the regulation of RET neuronal isoforms.

Here, we describe a novel function of AGO2 in the nuclear control of alternative polyadenylation, by using unbiased approaches and molecular tools. Our results indicate the involvement of AGO2 in alternative polyadenylation (APA) in neurons, including in control of the motor neuron-enriched GDNF receptor *Ret*.

## Results

### AGO2 is abundant in the nucleus of neuronal cells

To evaluate the nuclear localization of AGO2 in primary mouse motor neurons, we performed an immunofluorescence study, which revealed substantial AGO2 enrichment in motor neuron nuclei (**Figure 1A**). This unexpected abundance of AGO2 drove us to explore potential nuclear functions for AGO2. We have used a western blot (WB) analysis of a simple neuroblastoma cell line, NSC-34, and discovered that AGO2 was comparably abundant in nuclear and cytoplasmic fractions (**Figure 1B**). Accordingly, mass spectrometric (MS) profiling of cellular compartments, revealed that AGO2 abundance in the nucleus of NSC-34 cells, approximated cytoplasmic levels (**Figure 1C, Supplementary Table 4, Supplementary Figure 1A, B**). Therefore, AGO2 is particularly abundant in the nucleus of primary motor neurons and of a neuronal cell line.

**Figure 1.**
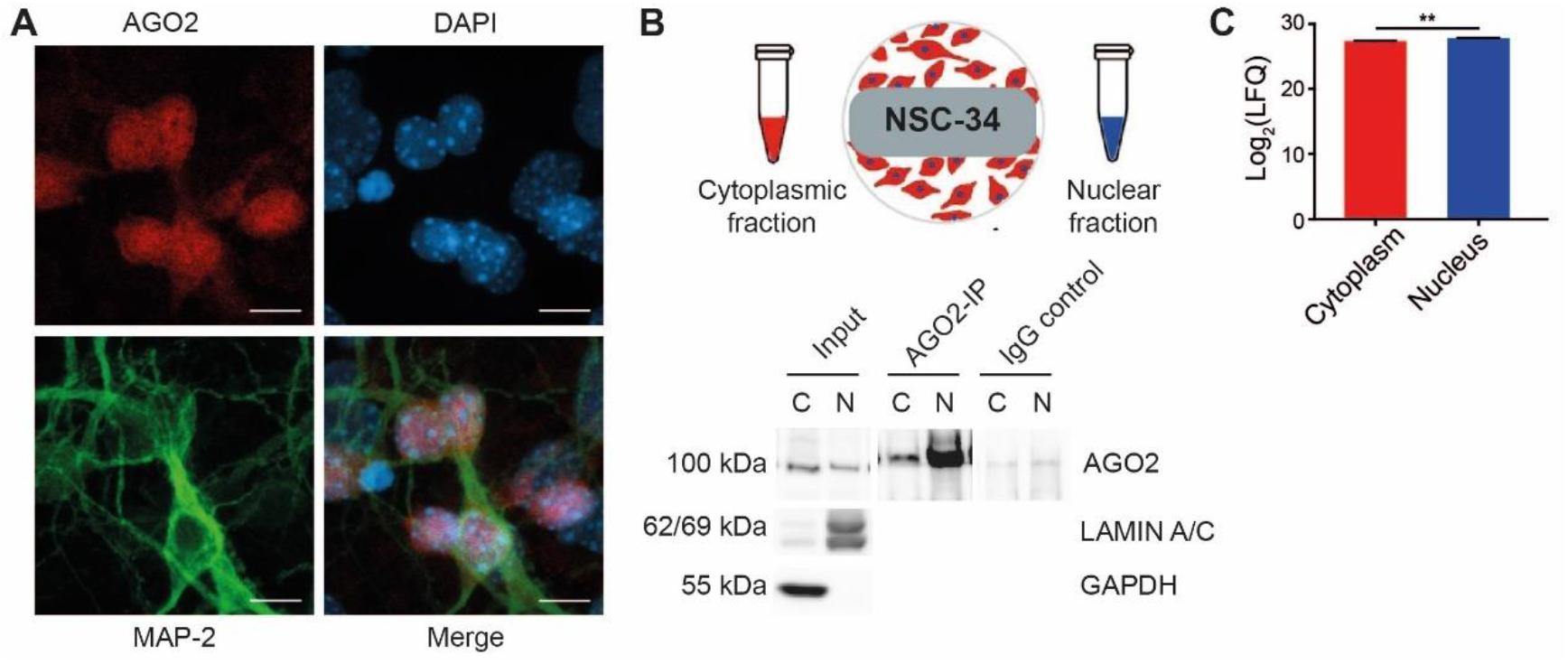
AGO2 is abundant in the nucleus of neuronal cells. **(A)** Micrographs of mouse primary motor neurons, stained with AGO2 antibody (red), neuron-specific microtubule associated protein MAP2 (green) and DAPI (blue). Scale bar – 10μm. Mouse primary motor neurons harvested on embryonic day E13.5 and cultured for 7 days before fixation and immunofluorescence analysis. **(B)** A diagram of NSC-34 subcellular fractionation protocol, and WB analysis of cytoplasmic ‘C’ or nuclear ‘N’ fractions. Input – lysates, or proteins co-immuno-precipitated with AGO2 or with IgG-control antibody. GAPDH/LAMIN A/C are cytoplasmic/nuclear markers, respectively. **(C**) Bar graph, depicting AGO2 mass spectrometry in cytoplasmic or nuclear fractions, identified by 24 unique peptides. *Student’s t-test* (S0=0.1) FDR p-value =0.00546.

### Identification of AGO2 nuclear interactors

To isolate nuclear AGO2-interacting proteins, we immunoprecipitated endogenous AGO2 (AGO2-IP) from nuclear NSC-34 fractions and analyzed by mass spectrometry. Data are available via ProteomeXchange with identifier PXD023112. We identified 472 AGO2-interacting proteins, which were enriched in nuclear AGO2-IP by at least two-fold, relative to non-specific IgG-control (q-value ≤0.05, using permutation-based false discovery rate (FDR)). Among AGO2 nuclear interactors, we report the three paralog co-factors TNRC6A, B and C, heterogeneous nuclear ribonucleoproteins (HNRNPs), nuclear paraspeckle components, ribosomal proteins, RNA helicases and RNA-splicing factors (**Figure 2A, Supplementary Table 5**). Interestingly, FUS, TDP-43, and HNRNPA2B1, whose mutated form is associated with motor neuron diseases also co-immunoprecipitated with AGO2.

**Figure 2.**
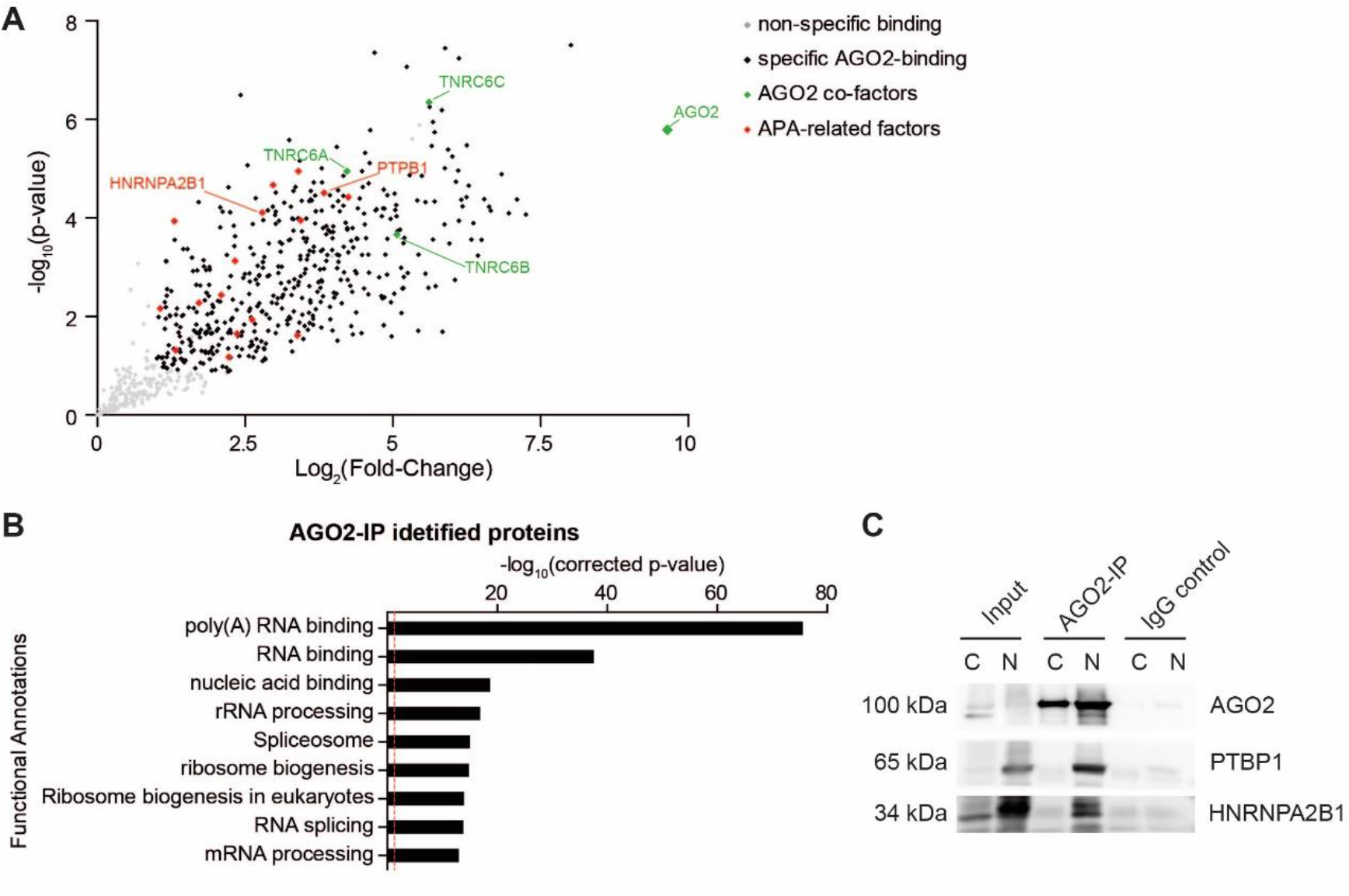
Identification of AGO2 interacting proteins in NSC-34 nuclei. **(A)** Scatter plot depicting proteins co-immunoprecipitated with AGO2 and identified by MS analysis. x-axis: log2 fold change of label-free peptide quantification values in AGO2-IP versus the IgG-control; y axis: log10of one-tailed *Student’s t-test* p-value. Grey-non-specifically bound proteins; Black-proteins specifically-bound by AGO2; Green TNRC6 A/B/C are AGO2 co-factors; Red – proteins with function in alternative polyadenylation. **(B)** Gene ontology analysis (DAVID 6.7 bioinformatic database, (Huang da et al., 2009)) of 472 nuclear AGO2 interacting proteins, shown as -log_10_(Bonferroni corrected p-value) of pathway enrichment (logarithmic scale). Dashed red line indicates a p-value of 0.05. **(C)** Western blot study of AGO2, PTBP1 and HnRNPA2B1, in cytoplasm ‘C’ and nucleus ‘N’ of NSC-34 after immunoprecipitation with antibody against AGO2 or non-specific IgG isotype control. Inputs are subcellular fraction lysates without immunoprecipitation.

Next, we tested if AGO2-interactors are identified in previously-characterized protein networks. The 472 AGO2 interactors are predicted by STRING (Franceschini et al., 2013) to create a dense protein-protein network, that is more prevalent than could be expected at random, when the full nuclear proteome (~2000 proteins) from the same cells is taken as background (p-value <1×10^-16^). Many nuclear AGO2 interactors are functionally annotated as RNA-binding proteins and involved in a variety of ribonucleoprotein complexes (DAVID Bioinformatics Resources 6.7 (Huang da et al., 2009), **Figure 2B**). Therefore, nuclear AGO2 in neuronal cells is associated with RNA-binding protein networks.

### AGO2 is involved in alternative polyadenylation

AGO2-nuclear interactors are enriched with proteins that are involved in alternative polyadenylation including direct APA factors CSTF1, CPSF7, NUDT21, PABPN1, RBBP6; and regulators PTBP1, PTBP2, PTBP3, ELAVL1, ELAVL2, ELAVL4, SRSF3, SRSF7, FUS, TDP-43 and HNRNPA2B1. This enrichment, that is more than can be expected at random, suggests potential functional association of AGO2 and the APA machinery (p-value = 0.0074 by hypergeometrical distribution test). We could also demonstrate the co-immunoprecipitation of at least two of the proteins with AGO2, PTBP1 and HNRNPA2B1, by WB analysis (**Figure 2C**).

To explore the hypothesis that AGO2 regulates APA, we performed next generation sequencing (NGS) of 3’ mRNA from NSC-34 cells, in which AGO2 was knocked-down by siRNA (**Supplementary Figure 2**). We computed molecular switches, as in (Rot et al., 2017), whereby a proximal (/distal) polyadenylation preference, is reciprocated by a new predominant polyadenylation signal that is positioned more distally (/proximally). A new proximal (/distal) APA site preference results in upregulation of shorter (/longer) 3’UTR, concomitant with downregulation of longer (/shorter) mRNA isoform (**Figure 3A, B**). We identified a total of 26,816 poly(A) sites, which were annotated to 11,157 mRNAs with at least one 3’ end isoform. Of these, 1952 pairs displayed proximal-over-distal polyadenylation preference switch in response to knockdown of AGO2, whereas 2933 pairs displayed a reverse distalover-proximal preference (**Supplementary Table 6**). A global transcriptome-wide change in polyadenylation preference was observed (Chi-squared test for the probability of a shift from proximal-to-distal preference or vice versa, p-value < 2.2E-16, **Figure 3C**), suggesting that APA regulation is a new and broad function of nuclear AGO2. Next, we focused our attention on 72 candidate transcripts, whose polyadenylation switches was the most significant after correction to multiple hypothesis (Benjamini-Hochberg based FDR adjusted p-value <0.05). Only nine Gencode annotated mRNAs (Frankish et al., 2019), displayed an AGO2-dependent polyadenylation switch and further harbor an AGO2-binding site at the vicinity (± 100bps) of the regulated poly-A site (AGO2-CLIP data from (Moore et al., 2015), **Table 1**).

**Figure 3.**
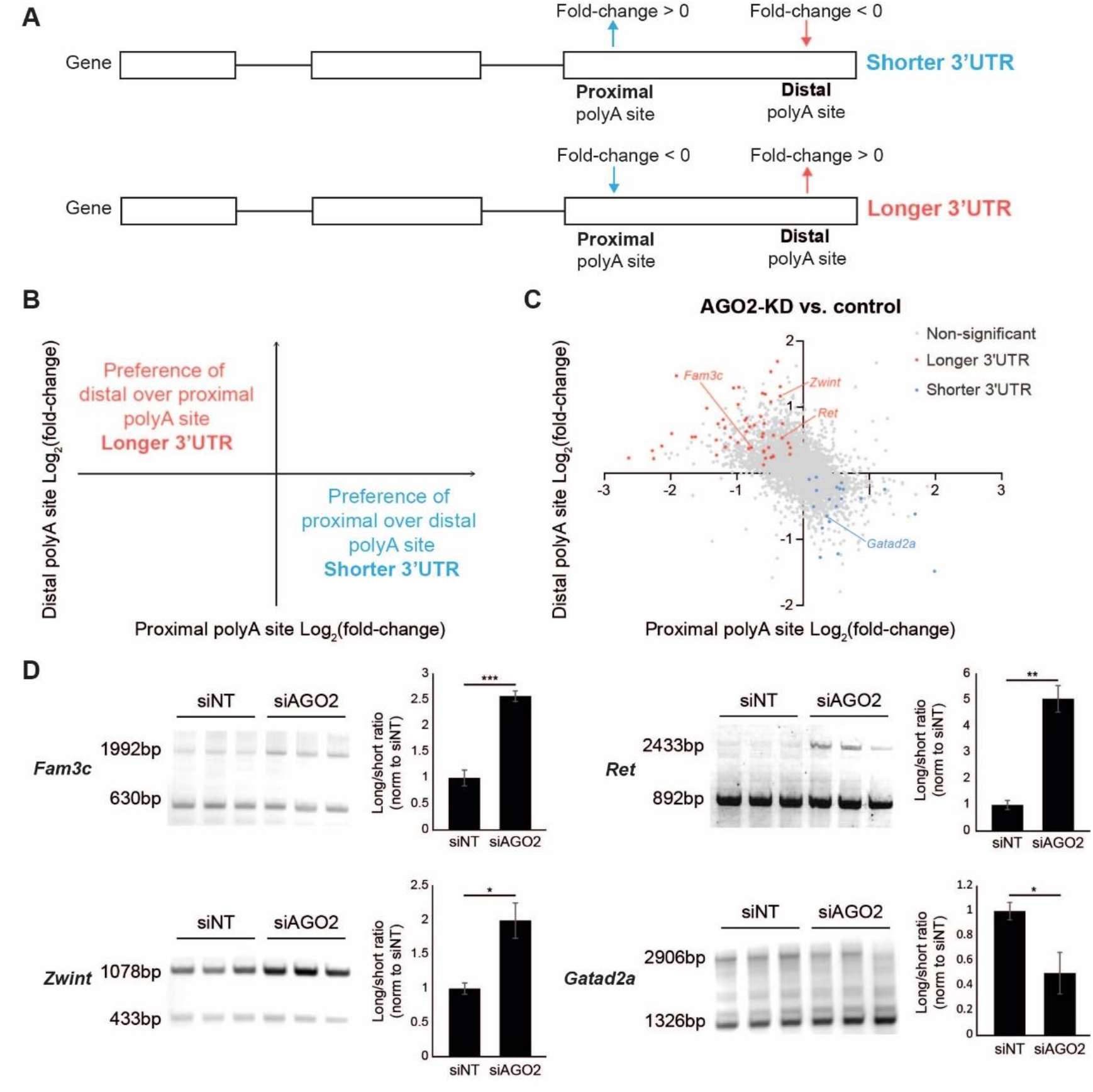
AGO2 activity controls alternative polyadenylation. Diagrams of **(A)** proximal to distal polyadenylation switch and **(B)** of data distribution over a scatter plot of proximal (x-axis) and distal (y-axis) polyadenylation ratio inferred from 3’ mRNA-sequencing. **(C)** Scatter plot of next generation RNA sequencing data, revealing polyadenylation ratio after AGO2 knockdown (siAGO2) vs non-targeting siRNA (siNT). mRNAs elongated/shorted their 3’UTR or were below statistical significance (red/ blue, grey respectively) by FDR-corrected p-values ≤0.05. **(D** Rapid amplification of cDNA ends (RACE), analysis of 3’ RNA polyadenylation preference by gel electrophoresis of RACE-PCR products and bar graph quantification. AGO2 knockdown (siAGO2) vs a non-specific siRNA control (siNT). Band densitometry normalized to controls. Average and SEM, two-tailed *Student’s t-test* p-value *≤0.05, **≤0.01, ***≤0.001. *Fam3c* – p-value = 0.00097; *Ret* – p-value = 0.0035; *Zwint* – p-value = 0.0217; *Gatad2a* – p-value = 0.05.

**Table 1.** AGO2-mediated alternative polyadenylation switches. Nine top hits exhibiting proximal/distal switch after knockdown of AGO2 with mRNA region wherein the PAS reside; Proximal/Distal FC – fold change of the usage of the chosen PAS between AGO2-KD and control; Proximal/Distal padj – adjusted p-value (correction for multiple hypothesis, FDR-based), for the AGO2-KD vs control comparison of the usage of the chosen PAS; UTR length difference – the difference (in base-pairs) between the proximal and distal PAS chosen as switch, +/− represent lengthening or shortening.

| Gene name | Proximal feature | Distal feature | Proximal FC | Distal FC | Proximal p-adj | Distal p-adj | UTR length difference |
| --- | --- | --- | --- | --- | --- | --- | --- |
| *Cxxc5* | utr3 | utr3 | 2.34 | 0.84 | 0.04496 | 0.03753 | -789 |
| *Fam3c* | utr3 | utr3 | 0.58 | 1.31 | 0.0223 | 0.00496 | +1369 |
| *Gatad2a* | utr3 | utr3 | 1.28 | 0.63 | 0.02205 | 0.00949 | -1581 |
| *Klc1* | utr3 | utr3 | 0.69 | 1.31 | 0.00019 | 0.00019 | +11818 |
| *Map6* | utr3 | utr3 | 0.66 | 1.26 | 0.00587 | 0.00178 | +16918 |
| *Pdlim7* | utr3 | exon | 0.59 | 1.65 | 0.00001 | 0 | +8360 |
| *Ret* | utr3 | utr3 | 0.80 | 1.44 | 0.01754 | 0.01793 | +2819 |
| *Trp53bp2* | utr3 | utr3 | 0.65 | 1.51 | 0.04462 | 0.00177 | 651 |
| *Zwint* | utr3 | utr3 | 0.787 | 2.24 | 0 | 0.00001 | 5798 |

We tested the polyadenylation switches of these 9 mRNAs by PCR-based 3’ rapid amplification of cDNA ends (3’ RACE; **Supplementary Figure 3**). This study revealed two polyadenylated isoforms for four mRNAs, out of the nine studied. Whereas AGO2 favors the proximal polyadenylation isoform of *Fam3c, Zwint* and *Ret*, and its knockdown resulted in preference towards the distal (longer) isoform, AGO2 activity contributes to distal *Gatad2a* polyadenylation (**Figure 3D**). Taken together, the involvement of AGO2 in APA regulation is evident via unbiased NGS study and analysis of specific targets.

### AGO2 mediates RET isoform expression via APA-regulation

Motor neuron-enriched *Ret* is a tyrosine-kinase receptor, activated by GDNF ligands (Airaksinen & Saarma, 2002; Arce et al., 1998; Baudet et al., 2008; Ibanez & Andressoo, 2017). In response to binding of GNDF, RET dimerizes and autophosphorylates several tyrosine residues at its intracellular domain, which docks downstream effectors (Airaksinen & Saarma, 2002; Ibanez & Andressoo, 2017). GDNF-RET signaling promotes motor neuron survival and is essential for neuromuscular junction development (Baudet et al., 2008; Zahavi et al., 2015).

We describe two *Ret* transcript variants, which correspond to the annotated Ret9 and Ret51 (Carter et al., 2001; Lee et al., 2003), with AGO2 binding sites (Moore et al., 2015), adjacent to *Ret* polyadenylation sequence (**Figure 4A**, blue).

**Figure 4.**
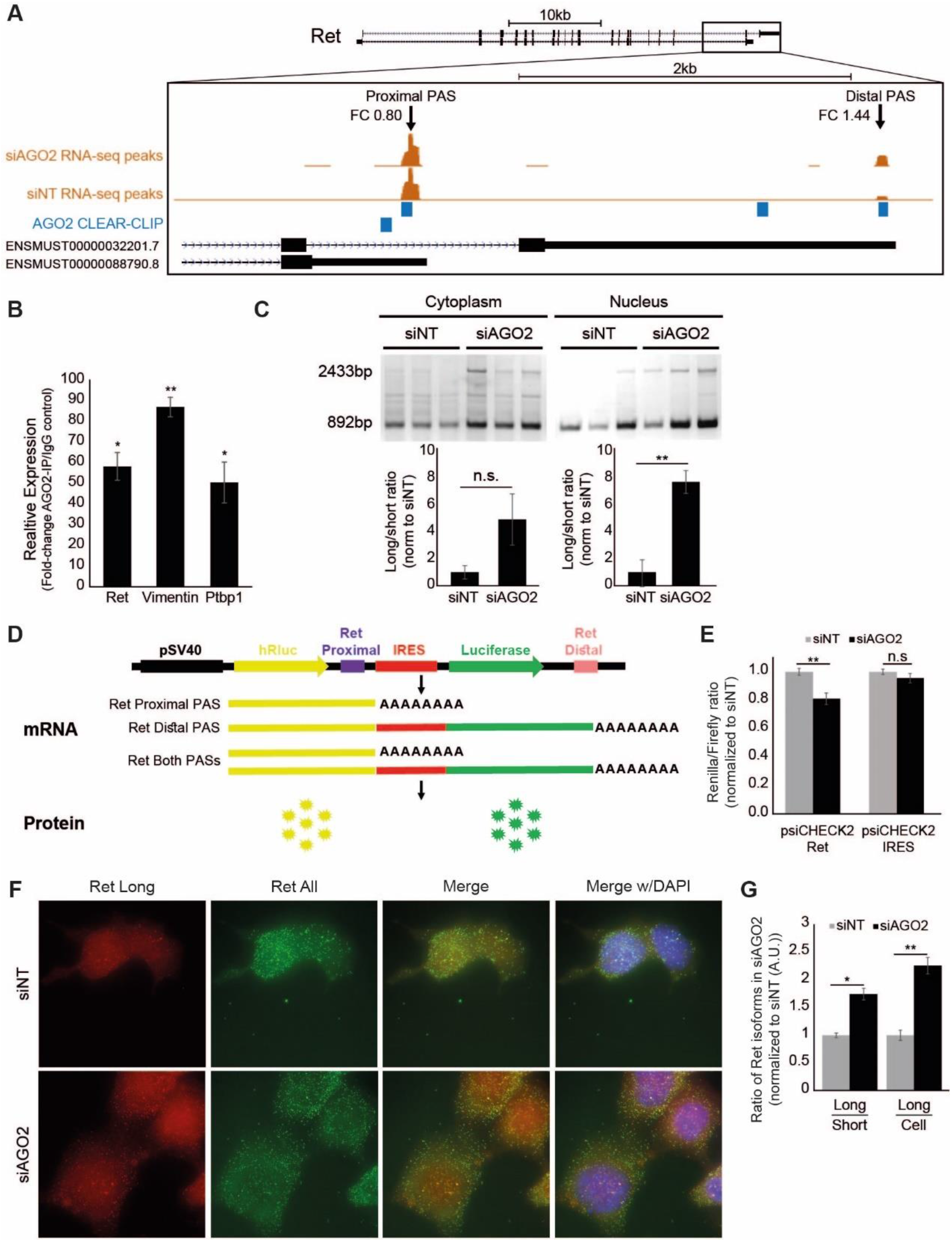
AGO2 controls Ret polyadenylation in a neuronal cell line. **(A)** Two annotated mRNA variants of the mouse *Ret* gene, depicted by 3’ next generation RNA sequencing reads at the proximal/distal polyadenylation site under basal conditions (siNT) or knockdown of AGO2 (siAGO2), with fold-change for each site (FC). The longer ENSMUST00000032201.7 encodes for Ret51 protein isoform, whereas the shorter, ENSMUST00000088790.8, encodes for Ret9. Position of AGO2 binding, at the 3’UTR of Ret mRNA, juxtaposed to the proximal polyadenylation site is based on CLEAR-CLIP data (Moore et al., 2015). **(B)** Real-time PCR quantification of mRNAs co-immunoprecipitated with AGO2 (AGO2-RIP). *Ret*, *Vimentin* and *Ptbp1* are associated with AGO2 in the nucleus of NSC-34 cells in a comparable manner. Average ± SEM, of data normalized to non-specific IgG isotype control. Two-tailed *Student’s t-test* p-value *≤0.05, **≤0.01, ***≤0.001. *Ret* - p-value = 0.01343; *Vimentin* - p-value = 0.00301; *Ptbp1* - p-value = 0.03803. **(C)** Gel electrophoresis of Ret mRNA isoforms in cytoplasm and nucleus, after AGO2 knockdown (siAGO2) and quantification of isoform densitometry ratio, normalized to control (siNT). Average ± SEM of 3 repeats, *two-tailed Student’s t-test*(p-value*≤0.05, **≤0.01, ***≤0.001). Cytoplasm – p-value = 0.11645; Nucleus – p-value = 0.00689. **(D)** Schematic diagram of the bicistronic reporter vector, adapted from (Deng et al., 2018). The vector contains Renilla luciferase protein ORF (hRluc) and firefly luciferase ORF (Luciferase) connected with one IRES, containing at the end of each ORF the proximal or distal PAS of Ret, correspondingly. The relative, activity of the proximal and distal Ret mRNA polyadenylation sites dictates the ratio between the two reporters. **(E)** AGO2 knockdown (siAGO2) resulted in preference of the distal polyadenylation site. Control reporter, lacking Ret polyadenylation sites (psiCHECK2-IRES), was unchanged. Average ± SEM of 5 repeats, *two-tailed Student’s t-test* (p-value*≤0.05, **≤0.01, ***≤0.001). psiCHECK2-Ret – p-value = 0.00378; psiCHECK2-IRES – p-value = 0.274293748. **(F)** Representative micrographs of Ret mRNA smFISH performed on NSC-34 cells previously subjected to AGO2-KD (siAGO2), or control (siNT) treatment 72hrs before fixation. smFISH was performed to detect only the long Ret variant (cy5, red) or both long and short Ret variants (Alexa 594, green). DAPI – nuclear DNA (blue). **(G)** Quantification of long and short Ret smFISH signal, 33 images per condition, 3 biological repeats. Average ±SEM, normalized to controls (siNT). Two-tailed Student’s t-test p-value for siAGO2 vs siNT. p-value * ≤0.05; **≤ 0.01, *** ≤0.001. Long/Short – p-value = 1.07394E-08; Long/Cell – p-value = 1.20772E-09.

To test if AGO2 directly binds *Ret* in the nucleus, we co-immuno-precipitated AGO2 with associated RNA (AGO2-RIP) from nuclear NSC-34 fraction. We demonstrated comparable levels of *Ret, Vimentin* and *Ptbp1* mRNAs, bound to AGO2 in the nucleoplasm, by quantitative PCR (qPCR, **Figure 4B**). Furthermore, AGO2-dependent isoform preference is evident in both nuclear and cytoplasm. Because mRNA undergoes directional nucleo-cytoplasmic export, the upregulation of Ret mRNA in the nucleus, in response to knocking down AGO2, is consistent with APA taking place in the nucleus (**Figure 4C**).

Next, we quantified the relative usage of two *Ret* APA *cis*-regulatory sequences in response to AGO2 levels, by using a bicistronic luciferase-reporter, similar to the one reported in (Deng et al., 2018). The reporter, which harbors the APA *cis*-regulatory sequences of *Ret* mRNA 3’, excludes many other potential regulatory elements that may reside on the transcript. The APA reporter assay in N2A cells, revealed that proximal Ret polyadenylation depends on AGO2 and that AGO2 knockdown resulted in preference of the distal *Ret* polyadenylation site (**Figure 4D, E**).

Finally, we used single-molecule fluorescent *in-situ* hybridization (smFISH) to test the cellular distribution of the *Ret* mRNA isoforms in NSC-34 cells and in mouse primary motor neurons (**Supplementary Figure 4A, B**). smFISH indicated the presence of both *Ret* mRNA variants in the soma and proximal neurites of the neurons. AGO2 knockdown resulted in a significant increase in total transcript copies and in upregulation of the long *Ret* mRNA variant mRNA, relative to all mRNA forms, (**Figure 4F, G**). Therefore, AGO2 regulates Ret mRNA alternative polyadenylation.

### GDNF signaling is implicated by AGO2-mediated Ret APA-regulation

*Ret* mRNA isoforms give rise to RET9 and RET51, which differ in their c-terminus (Ibanez, 2013; Rossel et al., 1997), and differentially activate downstream signaling pathways in response to GDNF (Hickey et al., 2009; Lian et al., 2017; Tsui-Pierchala et al., 2002).

To test whether AGO2-dependent control of *Ret* mRNA isoform switching is controlling the two protein isoforms, we performed WB analysis on lysates from NSC-34 cells, using an antibody that specifically binds to the longer RET51. We demonstrated that knockdown of AGO2 results in ~3-fold increase in the expression of RET51 (**Figure 5A, B**).

**Figure 5.**
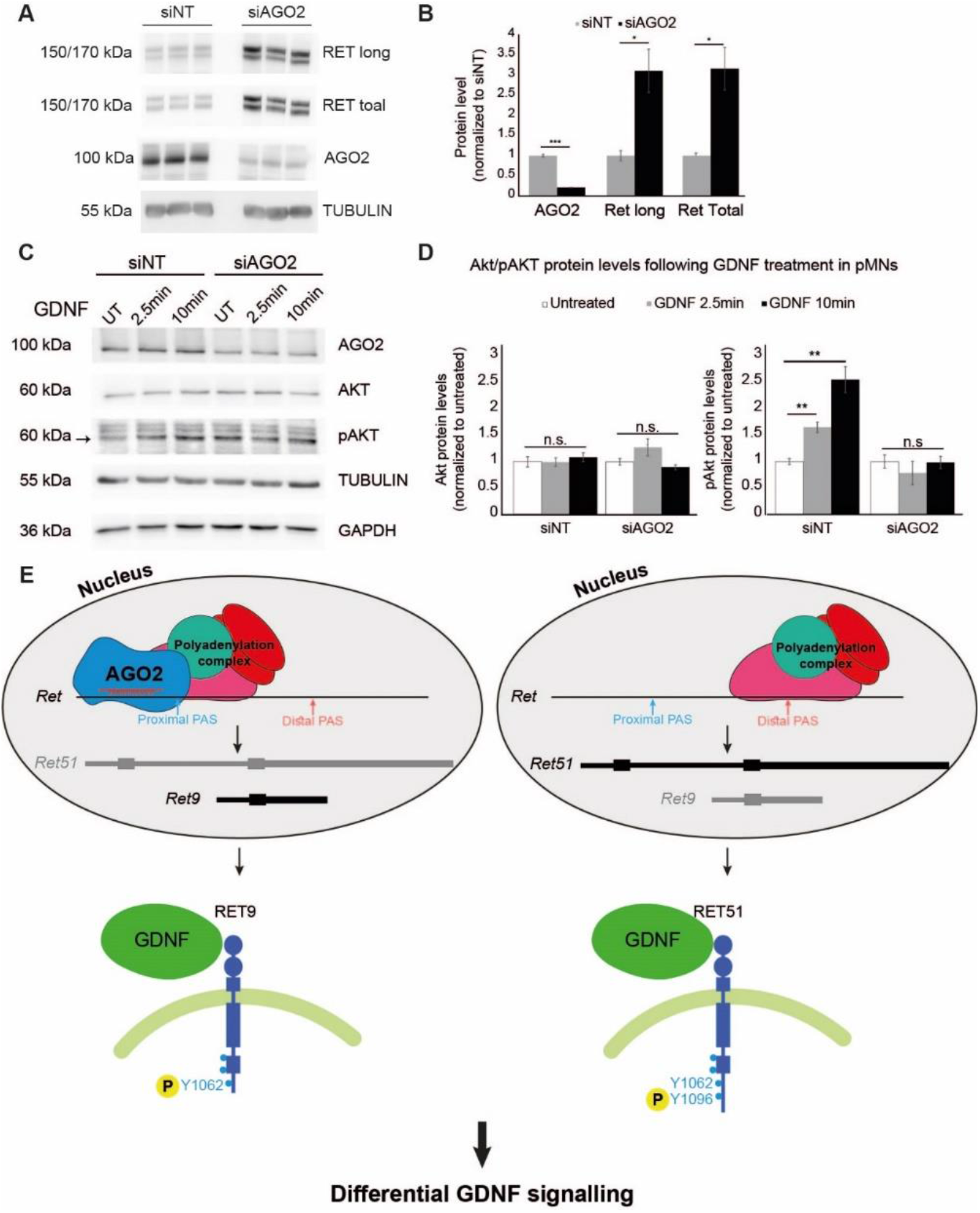
AGO2 controls Ret polyadenylation and GDNF signaling. **(A)** WB analysis and **(B)** band densitometry of AGO2 and RET protein isoforms, from NSC-34 cells, treated with siAGO2, normalized to Tubulin and to non-specific siRNA pool as control (siNT). Average ± SEM based on three biological repeats. Two-tailed Student’s t-test p-value * ≤0.05; *** ≤0.001. AGO2 – p-value = 2.05254E-05; RET long – p-value = 0.01888; RET total – p-value = 0.01512. **(C)** WB analysis of AGO2, AKT, pAKT, Tubulin and GAPDH in primary motor neurons, treated with GDNF for 2.5min or 10min. Representative image of three biological repeats. **(D)** Band densitometry of Akt and pAkt, normalized to Tubulin/GAPDH and to the untreated control. GDNF induces Akt phosphorylation. AGO knockdown (siAGO2 treatment) abolishes increased Akt phosphorylation. Average ± SEM based on three biological repeats. Two-tailed Student’s t-test p-value ** ≤0.01 p-values for AKT: siNT – GDNF 2.5min = 0.97736, GDNF 10min = 0.53249; siAGO2 – GDNF 2.5min = 0.19325, GDNF 10min = 0.25572. p-values for pAKT: siNT – GDNF 2.5min = 0.00445, GDNF 10min = 0.00361; siAGO2 – GDNF 2.5min = 0.4527, GDNF 10min = 0.92769. **(E)** A model for AGO2-mediated regulation of *Ret* polyadenylation in neurons. Nuclear AGO2 directs polyadenylation complex at the proximal poly(A) site of *Ret* mRNA, leading to the generation of RET9, whereas the distal polyadenylation site leads to the biogenesis of longer, RET51. RET9 and RET51 differs in signaling downstream of GDNF.

Next, we evaluated the changes in activation, of the GDNF-RET intracellular signaling, following AGO2 knockdown by following the phosphorylation state of AKT (pAKT) in mouse primary motor neurons (De Vita et al., 2000; Hayashi et al., 2000; Ibanez, 2013; Segouffin-Cariou & Billaud, 2000). WB analysis demonstrated an increase in pAKT in response to GDNF, relative to controls, and AGO2 knockdown resulted in dampening of AKT phosphorylation (**Figure 5C, D)**. Therefore, AGO2 regulates both RET isoform expression and the propagation of GDNF signal into pAKT. Together, we propose that AGO2-dependent control of alternative polyadenylation broadly impacts the neuronal transcriptome and specifically controls the balance between two translated isoforms of the GDNF receptor, RET, affecting intracellular signaling in motor neurons (**Figure 5E**).

## Discussion

In this study, we demonstrate that AGO2 is highly abundant in the nucleus of neurons and plays an unexpected role in broadly regulation of alternative polyadenylation. *Ret* mRNA, which encodes a tyrosine kinase receptor for GDNF, is probably among several mRNAs that are most sensitive to changes in AGO2 levels. Normally, the shorter RET9, is the predominant variant expressed in motor neurons (Lee et al., 2003). However, our data shows that AGO2 contributes to the preference of the short variant, *Ret9,* by actively controlling polyadenylation. Reduction in the expression of AGO2 changes the balance between the two *Ret* mRNA isoforms and attenuated intracellular signaling.

AGO2 is the main effector protein of miRNA-mediated silencing and more recently, nuclear AGO2 has been suggested to function in splicing and transcriptional regulation (Ameyar-Zazoua et al., 2012; Benhamed et al., 2012; Sarshad et al., 2018).

Alternative polyadenylation leads to the generation of 3’ mRNA variants (Di Giammartino et al., 2011; Elkon et al., 2013; Fu et al., 2018; Tian & Manley, 2017) that diversify the transcriptome expressed in the nervous system (Fontes et al., 2017; MacDonald, 2019; Miura et al., 2013; Ulitsky et al., 2012). APA contributes to differential mRNA localization (Mansfield & Keene, 2012; Taliaferro et al., 2016) and translation (Ainsley et al., 2014; Terenzio et al., 2018). Several RNA-binding proteins have been linked to APA regulation in the nervous system, including TDP-43 (Rot et al., 2017), FUS (Masuda et al., 2015; Schwartz et al., 2012), HnRNPA2/B1 (Martinez et al., 2016) and NOVA (Hwang et al., 2017; Licatalosi et al., 2008; Ule et al., 2003). Mutations in some of these APA regulators are associated with neurodegenerative or neuro-oncological diseases. In one such example, decreased levels of TDP-43, uncovers a cryptic polyadenylation site leading of STMN2 mRNA, which leads to a truncated non-functional STMN2 and eventually to the pathology of ALS (Melamed et al., 2019). Furthermore, global shortening of 3’UTRs was observed in ALS patients with C9ORF72 G4C2-repeat expansion (Prudencio et al., 2015), the most common genetic cause associated with ALS and frontotemporal dementia.

AGO2 associates with proteins that are directly involved in APA (CSTF1, CPSF2, CPSF3, NUDT21, PABPN1, RBBP6 and PTBP1), or regulate the process (ELAVL1, ELAVL2, SRSF3, SRSF7, FUS, TDP-43 and HNRNPA2B1) (Avendano-Vazquez et al., 2012; Elkon et al., 2013; Martinez et al., 2016; Rot et al., 2017; Tian & Manley, 2017). Accordingly, the global shift in polyadenylation preference in response to AGO2 manipulation, suggests that APA regulation is a novel broad function of nuclear AGO2.

RET, the receptor for the neurotrophic factor GDNF, is expressed in the developing and in the mature nervous systems, and is important for axonal growth and synapse development (Baudet et al., 2008; Bonanomi et al., 2012; Enomoto et al., 2001; Gould et al., 2008; Honma et al., 2010; Kramer et al., 2006; Pachnis et al., 1993; Tuttle et al., 2019), including the neuromuscular junction (Baudet et al., 2008). The formation of the short and long versions of RET protein, RET9 and RET51, seems to be the translational consequence of the two alternative mRNA isoforms, downstream of AGO2 activity. The balance between RET51 and RET9 determines the signaling output once activated downstream of GDNF (Besset et al., 2000; Coulpier et al., 2002; Crowder et al., 2004; Grimm et al., 2001; Hayashi et al., 2000; Lundgren et al., 2006; Perrinjaquet et al., 2010; Richardson et al., 2012; Segouffin-Cariou & Billaud, 2000; Tsui & Pierchala, 2010). We suggest that continuous AGO2 activity is required for maintaining RET9 predominance in motor neurons (Lee et al., 2003) which is needed for motor neuron survival (Airaksinen & Saarma, 2002).

It was previously shown that GDNF acts differently on motor neuron axons, facilitating growth and muscle innervation at axon terminals and survival pathways in the soma (Zahavi et al., 2015), which may be associated with differential *Ret* isoform distribution along the neuron.

Finally, the relevance of AGO2 to neurons and neurodegenerative diseases, including ALS, underscores the motivation to explore AGO2 neuronal functions but requires further studies to be elucidated.

Limitations: The abundance of AGO2 in the nucleus of neurons may be responsible for its involvement in APA. Therefore, we cannot rule out that in other cell types APA is also regulated by AGO. Furthermore, while direct AGO2 interactors are APA regulators, some of the effects may be indirect, for example by miRNA-based silencing of APA factor expression in the cytoplasm.

## Materials and Methods

### Cell culture and induction of cellular stress

NSC-34 cells (Cashman et al., 1992) were cultured in Dulbecco’s Modified Eagle Medium (DMEM, Biological Industries, 01-050-1A) supplemented with 10% Fetal Bovine Serum (FBS, Biological Industries, 04-001-1A), 1% penicillin-streptomycin (Pen-Strep, Biological Industries, 03-031-1B) and 1% L-glutamine (Biological Industries, 03-020-1B). Cells were grown at 37°C, 5% CO_2_. N2A cells were cultured in DMEM supplemented with 10% FBS and 1% Pen-Strep. Cells were grown at 37°C, 5% CO_2_.

### Culture of primary motor neurons

All experiments were performed in accordance with relevant guidelines and regulations of the Institutional Animal Care and Use Committee at Weizmann Institute of Science. Primary motor neurons (pMNs) were isolated and cultured as previously described (Milligan & Gifondorwa, 2011). Briefly, wild type (WT) ICR timed-pregnant females were sacrificed at mouse embryonic day 13.5 (E13.5), and spinal cords were dissected from embryos and dissociated enzymatically with papain (2mg/ml, Sigma, P4762). Motor neurons were separated over a gradient of Optiprep (Sigma, D1556) and plated on tissue culture plates pre-coated with 3μg/ml poly-Lysine (Sigma, P4707) and 3μg/ml laminin (Gibco, 23017015). Motor neurons were cultured with Neurobasal medium (Gibco, 211030-49) supplemented with 2% B27 (Gibco, 17504-044) 2% horse serum (Sigma), X1 Glutamax (Gibco, 35050-061), gentamycin 1 μg/ml (Sigma, G1272) and 1ng/ml CNTF (Peprotech, 450-50-25) and GDNF (Peprotech, 450-51-10). For GDNF treatment assay, cells were changed to starvation medium (Neaurobasal medium with antibiotics only) for 16hrs, then treated with 100ng/ml GDNF for 2.5min and 10min.

### Whole cell lysates and subcellular fractions

The protocol was adjusted to neuronal cells by harvesting cells using a low detergent-concentration lysis buffer supplemented with protease inhibitors and phosphatase inhibitors. One confluent 15cm NSC-34 culture dish was used per experimental repeat. Cells were washed ice-cold PBS prior to nuclear and cytosolic fractionation. All buffers used were prepared on the same day and supplemented with Protease Inhibitor Cocktail (Roche, 04693132001) and Phosphatase Inhibitor Cocktail (Roche, 04906837001). For WCL preparation, cells were harvested in ice-cold RIPA [50mM Tris-HCl, pH 7.5, 150mM NaCl, 1% NP-40, 0.5% deoxycholate (DOC), 0.1% SDS] and thoroughly mixed by vortex. The supernatant was collected after 10min incubation on ice, and 10min centrifugation at 15,871×g, 4°C. For nuclear and cytosolic fractionation, cells were harvested in ice-cold Lysis Buffer A [10mM Tris-HCl, pH 7.5, 10mM NaCl, 3mM MgCl2, 0.5% NP-40], incubated for 10 min on ice while vortexing every other minute. The cytoplasmic fraction was the collected supernatant after centrifugation for 10min at 15,871×g, 4°C. The nuclei were further pelleted, washed three times with ice-cold lysis buffer A by pipetting and then centrifuged for 10min at 15,871×g, 4°C. To gain the nuclear fraction, nuclei were resuspended in ice-cold RIPA, sonicated on ice at 30% power for three cycles of 10 sec with 20 sec intervals and centrifuged for 10min at 15,871×g, 4°C. The supernatant collected at this step was defined as the nuclear fraction. Protein concentrations were determined using Bio-Rad Protein Assay Dye Reagent (Bio-Rad, 500-0006). Fresh extracts were used for immunoprecipitation (IP) experiments, mass spectrometry (MS) or Western Blot (WB) analysis.

### Immunoprecipitation

50μl of Dynabeads Protein G (Novex by Life Technologies, 10004D) were mixed with 5μg of antibody (mouse-anti-AGO2; or mouse-IgG-isotype control), diluted in 200μl PBS + 0.1% TWEEN-20 (0.1% PBST) per sample, incubated by tilting at room temperature (RT) for 10min and washed three times with 0.1% PBST. Then, 300μl of protein extract (~1.5mg protein) was added to the bead-antibody complexes and incubated with rotation over-night (O.N.) at 4°C. For RNase treated samples, protein lysate was treated with RNase A/T1 (Thermo Scientific, EN0551, 18μg RNase A and 45U of RNase T1) for 30min at 37° with tilt prior to incubation with the antibody-beads complex. The following day, beads-antibody-antigen complexes were washed three times with PBS, resuspended in 150μl PBS and stored at −80°C until further analysis. For AGO2-RNA-Immunoprecipitation (AGO2-RIP), 25% of the precipitate was used for protein analysis, and 75% for RNA extraction, and was stored in 700μl QIAzol Lysis Reagent (Qiagen, 79306) until RNA purification step.

### Western blot

Cell extracts or IP-purified proteins were denatured by boiling (95°C) in X5 sample buffer (60mM Tris-HCl pH 6.8, 25% glycerol, 2% SDS, 14.4mM β-mercaptoethanol, 0.1% bromophenol blue) for 5min and resolved by 8%-10% SDS-PAGE, 100-120V, 70min. Proteins were transferred to a nitrocellulose membrane (Whatmann, 10401383) at 250mA, 70min. Membranes were stained with Ponceau (Sigma-Aldrich, P7170) to assess transfer quality, blocked for 1 hour at RT with 5% milk protein in PBS + 0.05% TWEEN-20 (0.05% PBST) and incubated, rocking, with primary antibodies O.N. at 4°C in Antibody-Solution [5% Bovine Serum Albumin, 0.02% sodium azide, 5 drops of phenol red in 0.05% PBST]. Following primary antibody incubation, membranes were washed three times for 5min at RT with 0.05% PBST and incubated for 1 hour at RT with horseradish peroxidase (HRP)-conjugated species-specific secondary antibodies. Membranes were washed three times for 5 min in 0.05% PBST at RT and protein bands were subsequently visualized by ImageQuant™ LAS 4000 (GE Healthcare Life Sciences) using EZ-ECL Chemiluminescence detection kit for HRP (Biological Industries, 20-500-120).

### Affinity Purification coupled to Mass Spectrometry proteomics (AP-MS)

Study design for the identification of AGO2-interacting proteins included four independent biological repeats, in each NSC-34 cell lysate from four confluent 10cm culture plates. Immunoprecipitated samples were subjected to AP-MS. Quantification was performed using MS1 based, label-free quantification (Shalit et al., 2015). The mass spectrometry proteomics data have been deposited to the ProteomeXchange Consortium (Deutsch et al., 2020) via the PRIDE (Perez-Riverol et al., 2019) partner repository with the dataset identifier PXD023112.

#### Sample preparation

For whole lysates samples were subjected to in-solution tryptic digestion using the suspension trapping (S-trap) as previously described (Elinger et al., 2019). Briefly, 50 ug of total protein was reduced with 5 mM dithiothreitol and alkylated with 10 mM iodoacetamide in the dark. Each sample was loaded onto S-Trap microcolumns (Protifi, USA) according to the manufacturer’s instructions. After loading, samples were washed with 90:10% methanol/50 mM ammonium bicarbonate. Samples were then digested with trypsin (1:50 trypsin/protein) for 1.5 h at 47°C. The digested peptides were eluted using 50 mM ammonium bicarbonate. Trypsin was added to this fraction and incubated overnight at 37°C. Two more elutions were made using 0.2% formic acid and 0.2% formic acid in 50% acetonitrile. The three elutions were pooled together and vacuum-centrifuged to dryness. Samples were kept at-80°C until further analysis.

For AGO2-IP, samples were subjected to on-bead tryptic digestion. Proteins were first reduced by incubation with dithiothreitol (5mM; Sigma-Aldrich) for 30 min at 60°C, and alkylated with 10 mM iodoacetamide (Sigma-Aldrich) in the dark for 30 min at 21°C. Proteins were then subjected to trypsin digestion (Promega; Madison, WI, USA) at trypsin:protein ratio of 1:50 at 37°C. Digestion was inhibited with trifluroacetic acid (1%) after 16 h, supernatant was isolated, desalted using solid-phase extraction columns (Oasis HLB, Waters, Milford, MA, USA) and stored in −80°C until further analysis.

#### Liquid chromatography

ULC/MS grade solvents were used for all chromatographic steps. For whole lysates, dry digested samples were dissolved in 97:3% H_2_O/acetonitrile + 0.1% formic acid. For all smaples (whole lysates and AGO2-IP), each sample was loaded using split-less nano-Ultra Performance Liquid Chromatography (10 kpsi nanoAcquity; Waters, Milford, MA, USA). The mobile phase was: A) H_2_O + 0.1% formic acid and B) acetonitrile + 0.1% formic acid. Desalting of the samples was performed online, using a Symmetry C18 reversed-phase trapping column (180μm internal diameter, 20mm length, 5μm particle size; Waters). The peptides were then separated using a T3 HSS nano-column (75μm internal diameter, 250mm length, 1.8μm particle size; Waters) at 0.35μL/min. Peptides were eluted from the column into the mass spectrometer using the following gradient: whole lysates – % to 20%B in 135 min, 20%-30% in 17 min, 30% to 90%B in 10 min, maintained at 90% for 5 min and then back to initial conditions; AGO2-IP – 4% to 30%B in 50 min, 30% to 90%B in 5 min, maintained at 90% for 5 min and then back to initial conditions.

#### Mass Spectrometry

For whole lysates – The nanoUPLC was coupled online through a nanoESI emitter (10 μm tip; New Objective; Woburn, MA, USA) to a Orbitrap Fusion Lumos mass spectrometer (Thermo Scientific) using a FlexIon nanospray apparatus (Proxeon). Data was acquired in data dependent acquisition (DDA) mode, using a 3-second cycle time method. MS1 resolution was set to 120,000 (at 200m/z) in the Orbitrap, mass range of 300-2000m/z, AGC of 4e5 and maximum injection time was set to 50msec. MS2 was performed in the ion trap, quadrupole isolation 1m/z, AGC of 1e4, dynamic exclusion of 30sec and maximum injection time of 50msec.

For AGO2-IP – The nanoUPLC was coupled online through a nanoESI emitter (10μm tip; New Objective; Woburn, MA, USA) to a quadruple orbitrap mass spectrometer (Q Exactive Plus, Thermo Fisher Scientific) using a FlexIon nanospray apparatus (Proxeon). Data was acquired in DDA mode, using a Top10 method. MS1 resolution was set to 70,000 (at 200m/z) and maximum injection time was set to 60msec, automatic gain control (AGC) target of 3e6. MS2 resolution was set to 17,500 and maximum injection time of 60msec, AGC target of 1e5. Quadrupole isolation window was set to 1.7m/z.

#### Raw Data processing

Raw data was processed with MaxQuant v1.6.0.16 (Cox & Mann, 2008), searched with the Andromeda against mouse (*mus musculus*) protein database as downloaded from Uniprot (www.uniprot.com) and appended with common lab protein contaminants. Enzyme specificity was set to trypsin and up to two missed cleavages were allowed. Fixed modification was set to carbamidomethylation of cysteines and variable modifications were set to oxidation of methionines, and deamidation of glutamines and asparagines. Peptide precursor ions were searched with a maximum mass deviation of 4.5 ppm and fragment ions with a maximum mass deviation of 20 ppm. Peptide and protein identifications were filtered at an FDR of 1% using the decoy database strategy (MaxQuant’s “Revert” module). The minimal peptide length was 7 amino-acids and the minimum Andromeda score for modified peptides was 40. Peptide identifications were propagated across samples using the match-between-runs option checked. Searches were performed with the label-free quantification (LFQ) option selected.

#### Proteomics Statistical Analysis

ProteinGroups output table was imported from MaxQuant to Perseus environment v1.6.0.2 (Tyanova et al., 2016). Quality control excluded reverse proteins, proteins identified by a single peptide, and contaminants. For lysate input analysis, quantitative comparisons were calculated based on log2-transformed LFQ values. Protein groups required ≥3 valid values/group. Missing data were replaced using imputation, assuming normal distribution with a downshift of 1.6 standard deviations and a width of 0.4 of the original ratio distribution. *Student’s t-test* with S0=0.1 was performed with FDR p-value≤0.05 for pairs of cytoplasmic fraction and nuclear fraction samples in each condition.

For AGO2-pull down analysis, non-specific IgG-isotype control binders were excluded by log2-transformed LFQ values. Protein groups required ≥3 valid values/group. Missing data were replaced using imputation, assuming normal distribution with a downshift of 1.6 standard deviations and a width of 0.4 of the original ratio distribution. Enriched AGO2 interactors were called by *Student’s t-test* (AGO2-IP vs corresponding IgG-control) with S0=0.1 and FDR p-value≤0.05 and fold-change threshold of 2-fold enrichment.-

### Immunofluorescence

NSC-34 cells were seeded at a density of 75,000cells/cm^2^ on 13mm glass cover-slips (Thermo Fisher Scientific), precoated with 0.002% poly-L-lysine (Sigma-Aldrich, P4707) and cultured for 2 days at 37°C and 5% CO_2_. pMNs were seeded 200,000 cells per coverslip, precoated as described. Cells were then washed with PBS, fixed with 4% paraformaldehyde (PFA) for 15min at RT, permeabilized with PBS containing 0.2% (vol/vol) Triton X-100 and blocked using CAS block (Invitrogen, 008-120) for 10min at RT. After incubation with primary antibodies O.N., 4°C, cells underwent three washes with PBS (5min each) and incubated for 1 hour at RT with secondary antibodies conjugated with Cy2, Cy3 or Cy5 diluted in CAS block. Glass cover slips were mounted on Superfrost microscope slides (Thermo Fisher Scientific) with Fluoroshield mounting media containing DAPI (Sigma-Aldrich, F6057). Fluorescence images were captured using a Zeiss LSM780/800 Laser Scanning confocal microscope system.

### siRNA knockdown

Dharmacon siGenome SMARTpool siRNAs against mouse AGO2 (M-058989-01-0005) was used at a final concentration of 20nM to knockdown AGO2 in NSC-34 cells, N2A cells or primary MNs. siRNAs were transfected into NSC-34 or N2A cells using Lipofectamine RNAiMAX Reagent (Thermo Fisher Scientific, 13778-075) according to the manufacturer’s instructions. For pMNs, Dharmafect 4 (Dharmacon, T-2004-02) reagent was used for siRNA transfection. Dharmacon siGENOME Non-targeting pool #2 (siNT, D-001206-14-05) were used as controls. Knockdown efficiency was assessed by RNA extraction followed by quantitative Real-time PCR (qRT-PCR).

### Quantitative Real-time PCR

Total RNA from cultured NSC-34 cells or primary motor neurons was isolated using DirectZole RNA Purification Kit (Zymo Research, R2052), and reverse transcribed using qScript cDNA Synthesis Kit (Quanta Biosciences, 95047-100). Quantitative realtime PCR was performed using StepOnePlus real-time PCR instrument (Applied Biosystems), in >3 independent biological repeats and technical duplicates. KAPA SYBR FAST ABI Prism (KAPA Biosystems, KK4604) was used for detection of mRNAs. *Gapdh* or *TBP* were used as reference genes for normalization of expression levels. Statistical analysis was performed using *Student’s t-test.* For AGO2-RIP experiments, RNA was extracted using miRNeasy micro Kit (Qiagen, 217084), and reverse transcribed using the miScript II RT Kit (Qiagen, 218161). Quantitative realtime PCR was performed using StepOnePlus real-time PCR instrument, in >3 independent biological repeats and technical duplicates. miScript SYBR Green PCR (Qiagen, 218073) and KAPA SYBR FAST ABI Prism kits were used for detection of miRNAs and mRNAs respectively. *U6* and *TBP* were used as reference genes for normalization of miRNA and mRNA levels, respectively. All primer sequences are described in Supplementary Table 1.

### 3’-RNA sequencing

#### cDNA Libraries Preparation

The poly(A)seq cDNA libraries were generated using the reverse QuantSeq 3’ mRNA-seq Library Prep Kit for Illumina (Lexogen). Libraries were prepared from 400ng of total RNA according to kit instructions. Single-end sequencing (60bp) was performed on Illumina HiSeq2500 with a Rapid Run flow-cell. RNA-seq data are available in the ArrayExpress (Athar et al., 2019) database (http://www.ebi.ac.uk/arrayexpress) under accession number E-MTAB-9907.

#### Data Processing

Bioinformatic analysis pipeline was performed as described in (Rot et al., 2017). Briefly, all data sets were processed by aligning the reads to the reference mouse genome (mm10) using STAR aligner (Dobin et al., 2013) with default parameters and with the GTF annotations from Ensembl. The polyadenylation events were determined by tagging only one position per alignment (the first 5’ aligned nucleotide). Alignments containing stretches of six consecutive A or with 70% A coverage in any 10-nt sub-window in the region [-10..10] surrounding the polyadenylation events were filtered out. Polyadenylation events were ranked by read count in descending order and only the high-ranking events that are more than 125nt apart were considered as dominant poly(A) sites and therefore in the analysis. This resulted in the global poly(A) site database. To allow variation in cleavage precision, the per-experiment expression of each poly(A) site was computed by summing the read counts that identify any position in the region [-5..5] surrounding the polyA cleavage sites.

#### Statistical Analysis

DEXSeq algorithm was applied to identify regulated poly(A) sites in genes. Count values of all poly(A) sites remaining after filtering for all replicates for control and AGO2-KD 3’end sequencing experiments were input. The output was foldchange (log2) and FDR adjusted p-value for each site. Genes in which no poly(A) site reached significance (p≤0.05) were classified as controls. In regulated genes with more than one poly(A) site, only two significantly changed sites (adjusted p-value ≤0.05) with highest difference in fold change were selected for each gene, additionally requiring that fold changes are of opposite direction. If a proximal site had fold change <0, the site was marked as repressed, and if fold change >0, marked as enhanced (the reverse holds for distal sites). In control genes, the two poly(A) sites with highest read count were considered for further analysis; the proximal and distal control poly(A) sites were further labeled as control-down and control-up (dependent on their fold change). The poly(A) site pairs were further classified into different types of alternative polyadenylation (same exon, composite exon, and skipped exon (Rot et al., 2017)), using the gene level annotation which was computed by linearizing the Ensembl gene annotation by merging the transcript annotation.

### Semi-quantitative PCR-based Alternative polyadenylation assay (3’RACE-like)

RNA was extracted from the cells using DirectZole RNA MiniPrep (Zymo Reseach, R2052) according to kit instructions. cDNA was produced from 500ng of RNA using the high-capacity reverse transcription kit (Applied Biosystems, 4374966), and Oligo(dT)-VN primers with an adapter sequence according to the manufacturers’ instructions. To evaluate alternative polyadenylation (APA) events, a semi-quantitative RT-PCR reaction was performed using Q5 Hot Start High-Fidelity DNA polymerase (NEB, M0493S) in a LabCycler thermocycler (SensoQuest). 5ng cDNA was used for PCR and amplified using 29 cycles. RT-PCR products were separated on 1.5% agarose gel and visualized using MiniBIS Pro (DNR Bio-Imaging Systems). Quantification of APA band intensities were determined using ImageJ software and an average of three biological replicates was plotted. Forward primers for each gene were designed to be in the last exon or one upstream. Reverse primer was designed for the adapter sequence used in the cDNA generation step. All primers are listed in Supplementary Table 1.

### Single-molecule Fluorescent in-situ Hybridization (smFISH)

Probe library construction, hybridization procedure and imaging conditions were previously described (Farack et al., 2019; Itzkovitz & van Oudenaarden, 2011; Lyubimova et al., 2013; Raj et al., 2008). Probe libraries were designed using the Stellaris FISH Probe Designer (Biosearch Technologies) and consisted of 41-48 probes each of length 20 bps, complementary to the mRNA sequence of either both *Ret* variants (Ret-all, *Ret9, Ret51),* or a unique sequence in the long *Ret* variant (Ret-long, *Ret51).* Ret-all probe library was coupled to Alexa594 (CAL Fluor Red 610, Stellaris, Biosearch Technologies); Ret-long probe library was coupled to Cy5 (Quasar 670, Stellaris Biosearch Technologies). NSC-34 cells were seeded 500,000 cells per coverslip on 22×22mm glass coverslips (Thermo Fisher Scientific), coated with 0.002% poly-L-lysine (Sigma-Aldrich, P4707). pMNs were seeded 500,000 cells per coverslip, precoated as described. Cells were transfected with siAGO2 or siNT (as described) and cultured for 72hrs at 37°C and 5% CO_2_. Cells were then washed with nuclease-free PBS, fixed with 4% paraformaldehyde (PFA) for 10min at RT, washed again with PBS and incubated in 70% ethanol for 1-3 days at 4°C. Formamide concentration of the washing and hybridization buffers was 25%. Fixed cells were washed twice in 2xSSC, then incubated for 1.5hrs with the washing buffer before hybridizing with the probe libraries. Probes were diluted in hybridization buffer and cells were incubated O.N. at 30°C. The following day, cells were washed twice with washing buffer for 30min at 30°C. Next, cells were washed with GLOX buffer (2xSSC, 10% glucose, 10mM Tris) for 5min at RT. Nuclei were stained with DAPI (Sigma-Aldrich, D9542), then washed again with GLOX buffer twice, 5min each wash. Slides were mounted using ProLong Gold (Molecular Probes, P36934).

#### Imaging

smFISH imaging was performed using a Nikon-Ti2 Eclipse inverted fluorescence microscope equipped with a 100×oil-immersion objective and a Photometrics iXon Ultra 888 EMCCD camera using NIS-Elements Advance Research software (Nikon Instruments Inc.). Quantification was performed on stacks of optical sections with Z spacing of 0.3 μm.

#### Image Analysis, quantification and statistics

Imaris Cell Imaging Software 9.2.1 (Oxford Instruments Group), was used to analyze images, including detection and quantification of smFISH signal of each probe and colocalization of the signals from both probes. Cell number for each image was calculated based on DAPI signal using ImageJ v1.52n (Schindelin et al., 2012) software for per-cell quantification. Images were visualized and processed using ImageJ v1.52. Statistical analysis was performed using *Student’s t-test*.

### Cloning of APA reporter vector

Construction of the Luciferase-based APA reporter vector was performed based on (Deng et al., 2018) using Restriction-Free Cloning (Unger et al., 2010) upon the psiCHECK-2 dual luciferase assay vector. All amplification reactions were performed using Q5 Hot Start High-Fidelity DNA polymerase (NEB, M0493S) in a LabCycler thermocycler (SensoQuest). The HSV-TK promoter was replaced with IRES sequence to allow the transcription of a bicistronic mRNA. The vector created following this step was used as control (psiCHECK-2-IRES). For evaluation of Ret APA (psiCHECK-2-Ret), the generic poly(A) signal at the end of the first ORF was replaced with Ret proximal poly(A) site and with Ret distal poly(A) site for the second ORF (cleavage site +/− 200bps for each poly(A) site). All vectors were verified by sequencing. Primers used for cloning are listed in Supplementary Table 2.

### Dual Luciferase Activity Assay Hela

N2A cells were co-transfected using Lipofectamine 2000 Transfection Reagent (Thermo Fisher Scientific, 11668019) with either the control luciferase vector (psiCHECK-2-IRES) or Ret construct (psiCHECK-2-Ret), combined with either siAGO2 or siNT. Cells were harvested 72hrs post-transfection. The resulting lysates were used to measure the humanized renilla (hRluc) and firefly (hluc) luciferase activities with the Dual-Luciferase Reporter Assay kit (Promega, E1960) according to the manufacturer’s instructions. Luminesce was read using Veritas luminometer (Turner BioSystems, CA, USA). Statistical analysis was performed using *Student’s t-test* for 5 replicates for each condition.

### Antibodies

All Antibodies used in this study are listed in Supplementary Table 3.

## Acknowledgements

E.H. is the Mondry Family Professorial Chair and head of the Nella and Leon Benoziyo Center for Neurological Diseases at Weizmann Institute of Science. We thank Topaz Altman and Eran Perlson (Tel Aviv University) for discussions and reagents, members of the Hornstein lab for helpful critiques. Research in the Hornstein lab was supported by the Radala Foundation the Minerva Foundation with funding from the Federal German Ministry for Education and Research, ISF Legacy grant 828/ 17; Target ALS (118945); the European Research Council European Union’s Seventh Framework Programme (FP7/2007-2013)/ERC grant agreement 617351; Israel Science Foundation (135/16, 392/21); the ALS Therapy Alliance; AFM-Te’ le’ thon (20576); the Motor Neurone Disease Association (UK); The Thierry Latran Foundation for ALS Research; ERA-Net Research Programme on Rare Diseases (FP7); Yeda-Sela, Yeda-CEO; the Israel Ministry of Trade and Industry; Y. Leon Benoziyo Institute for Molecular Medicine; the Benoziyo Center Neurological Disease; the Kekst Family Institute for Medical Genetics; the David and Fela Shapell Family Center for Genetic Disorders Research; the Crown Human Genome Center; the Nathan, Shirley, Philip, and Charlene Vener New Scientist Fund; the Julius and Ray Charlestein Foundation; the Fraida Foundation; the Wolfson Family Charitable Trust; the Abney Foundation; Merck; Maria Halphen; and the estates of Fannie Sherr, Lola Asseof, Lilly Fulop, and. Edward and Janie Moravitz.

## Declaration of Interests

The authors declare no competing interests.

## Supplementary Figures

**Supplementary Figure 1.**
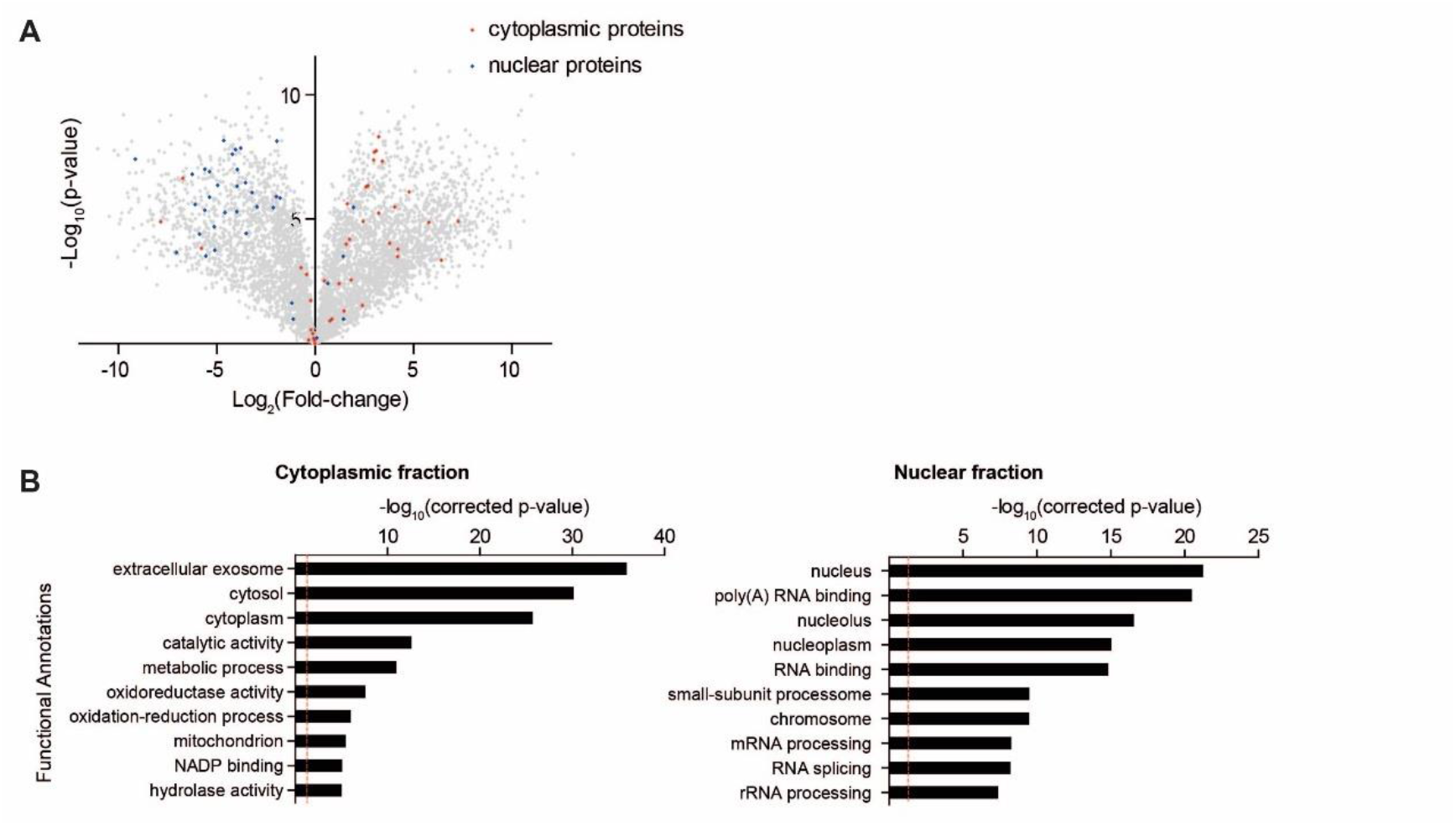
A platform for characterizing AGO2 protein-interactions in neuronal nuclei. **(A)** Volcano plot depicting the distribution of all proteins identified by MS analysis of cytoplasmic and nuclear fractions lysates. x-axis: log2 fold change of label-free peptide quantification values in cytoplasmic fraction versus nuclear fraction; y axis: log10 of two-tailed *Student’s t-test* p-value. Grey-all proteins; Red/Blue-cytoplasmic/nuclear marker proteins obtained from protein atlas (Thul et al., 2017). **(B)** Gene ontology analysis of proteins that are enriched in either nuclear (top panel) or cytoplasmic (lower panel) fractions (>100 fold), shown as -log10(Bonferroni corrected p-value) of pathway enrichment (logarithmic scale). Dashed red line indicates a p-value of 0.05. Analysis performed with DAVID 6.7 bioinformatic database(Huang da et al., 2009). Categories enriched in each fraction provide further support of efficient nuclear-cytoplasmic separation.

**Supplementary Figure 2.**
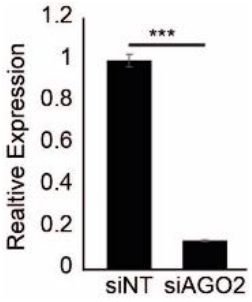
Quantitative Validation of AGO2-KD using siAGO2. Real-time PCR quantification of *Ago2* mRNA in NSC-34 cells treated with siAGO2 vs siNon-Targeting (siNT) as control. Average ± SEM of 3 repeats, data normalized to siNT. Two-tailed *Student’s t-test* p-value *≤0.05, **≤0.01, ***≤0.001; p-value = 0.000013854.

**Supplementary Figure 3.**
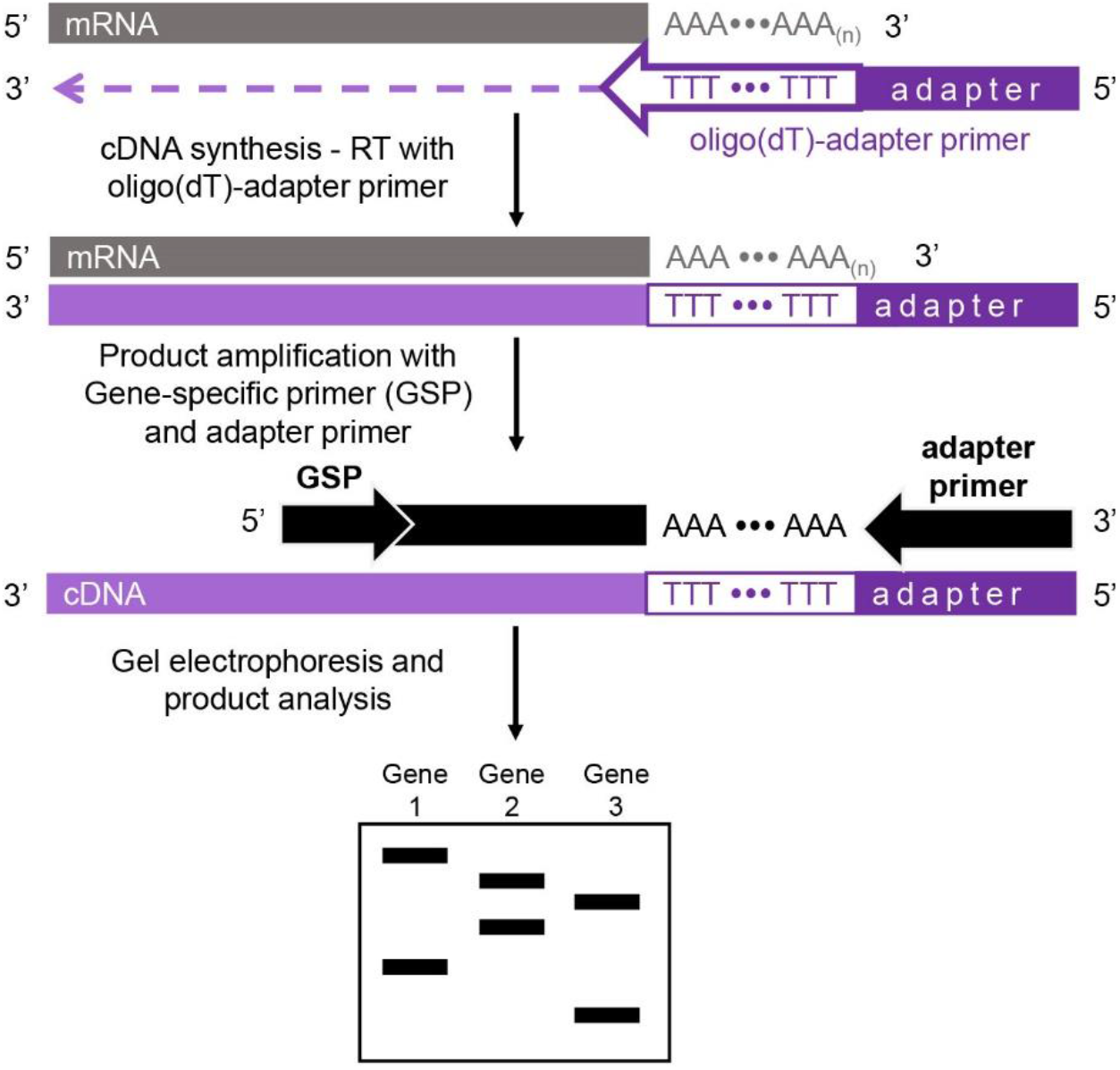
3’RACE-like PCR demonstration of APA. An oligo(dT)-VN primer mix (V-any nucleotide but T, N-any nucleotide) planned to capture transcript 3’ end was used to generate cDNA with an adapter sequence subsequently downstream of the poly(A) tail. Gene-specific PCR reaction was then performed and analyzed by gel electrophoresis depict differential usage of PAS, and hence different poly(A) tail length.

**Supplementary Figure 4.**
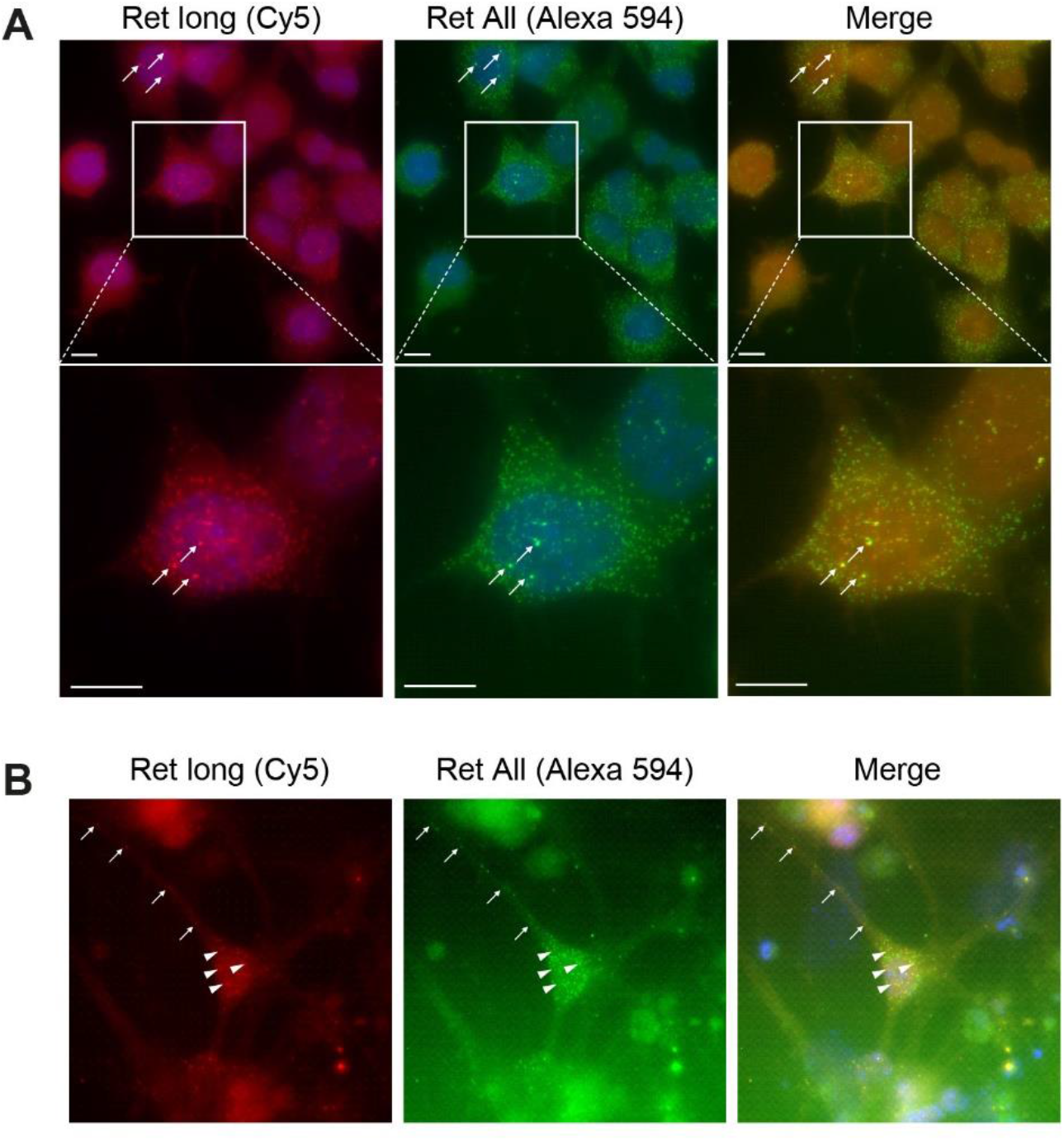
*Ret* APA isoforms smFISH study in mouse neurons. smFISH probes detect only the long variant (cy5, red) or both long and short variants (Alexa 594, green) of Ret. Arrowheads and arrows depict the long variant in NSC-34 cells **(A)** and in soma and neurites, respectively, in mouse primary motor neurons **(B)**. scale bar – 10μM.

## Notes

### Competing Interest Statement

The authors have declared no competing interest.

